# Control of mechanical pain hypersensitivity through ligand-targeted photoablation of TrkB positive sensory neurons

**DOI:** 10.1101/233452

**Authors:** Rahul Dhandapani, Cynthia Mary Arokiaraj, Francisco J. Taberner, Paola Pacifico, Sruthi Raja, Linda Nocchi, Carla Portulano, Federica Franciosa, Mariano Maffei, Ahmad Fawzi Hussain, Fernanda de Castro Reis, Luc Reymond, Emerald Perlas, Simone Garcovich, Stefan Barth, Kai Johnsson, Stefan G. Lechner, Paul A. Heppenstall

**Affiliations:** EMBL Mouse Biology Unit, Via Ramarini 32, Monterotondo 00015, Italy.; Molecular Medicine Partnership Unit (MMPU), Heidelberg, Germany.; Institute of Pharmacology, Heidelberg University, Im Neuenheimer Feld 366, 69120 Heidelberg, Germany; Department of Gynecology and Obstetrics, University Hospital RWTH Aachen, Pauwelsstrasse 30, 52074, Aachen, Germany.; Ecole Polytechnique Federale de Lausanne, Institute of Chemical Sciences and Engineering (ISIC), Institute of Bioengineering, National Centre of Competence in Research (NCCR) in Chemical Biology, 1015 Lausanne, Switzerland; Institute of Dermatology, Catholic University of the Sacred Heart, Largo A. Gemelli 8, 00168, Rome, Italy; South African Research Chair in Cancer Biotechnology, Institute of Infectious Disease and Molecular Medicine (IDM), Department of Integrative Biomedical Sciences, Faculty of Health Sciences, University of Cape Town, Anzio Road, Observatory 7925, South Africa.

**Keywords:** Mechanical allodynia, Neuropathic pain, Low-threshold mechanoreceptors, TrkB, primary afferents, ablation, optogenetics, photoablation, animal behaviour

## Abstract

**Summary:** Mechanical allodynia is a major symptom of neuropathic pain whereby innocuous touch evokes severe pain. Here we identify a population of peripheral sensory neurons expressing TrkB that are both necessary and sufficient for producing pain from light touch after nerve injury. Mice in which TrkB-Cre expressing neurons are ablated are less sensitive to the lightest touch under basal conditions, and fail to develop mechanical allodynia in a model of neuropathic pain. Moreover, selective optogenetic activation of these neurons after nerve injury evokes marked nociceptive behavior. Using a phototherapeutic approach based upon BDNF, the ligand for TrkB, we perform molecule-guided laser ablation of these neurons and achieve long-term retraction of TrkB positive neurons from the skin and pronounced reversal of mechanical allodynia across multiple types of neuropathic pain. Thus we identify the peripheral neurons which transmit pain from light touch and uncover a novel pharmacological strategy for its treatment.

**Highlights:**

- TrkB^+^ neurons detect light touch under basal conditions
- TrkB^+^ neurons convey mechanical allodynia in neuropathic pain states
- A photosensitizing derivative of BDNF allows for photoablation of TrkB^+^ neurons
- BDNF-guided photoablation reverses allodynia in multiple types of neuropathic pain

## Introduction

In neuropathic pain patients, hypersensitivity to light touch can develop to the extent that movement of a single hair shaft is sufficient to provoke severe pain (Treede et al., 1992). This is difficult to treat using conventional analgesics such as opioid or NSAIDS, and it impacts greatly upon quality of life due to the pervasive nature of mechanical stimuli; for example, small movements of the body, or the weight of clothing can cause severe pain in neuropathic patients (Koltzenburg, 2000). While much recent progress has been made in delineating the spinal circuits that gate mechanical pain (Duan et al., 2014; Foster et al., 2015; Peirs et al., 2015; Torsney and MacDermott, 2006), the identity of the sensory neurons that input this sensation into the spinal cord is less clear (Arcourt and Lechner, 2015).

Hypothetically, mechanical hypersensitivity could be mediated either by sensitization of nociceptors (hyperalgesia), or through integration of input from low threshold mechanoreceptors into pain transmitting circuits (allodynia) (Treede et al., 1992). In human studies, there is little evidence for nociceptor sensitization, and most reports indicate that mechanical allodynia is conveyed by myelinated A-fibre mechanoreceptors, although it is not known which subtype (Koltzenburg, 2000). Thus, differential block of these nerves alleviates brush evoked pain (Campbell et al., 1988), and the short latency of pain perception is indicative of the fast conduction velocity of A-fibers (LaMotte et al., 1991). In experimental animal studies the situation is less clear. For example, it has been demonstrated that mice develop mechanical allodynia in neuropathic pain models even when all nociceptors are genetically ablated (Abrahamsen et al., 2008). In contrast, Isolectin B4 positive nociceptors have also been implicated as drivers of allodynia (Tarpley et al., 2004). Another study proposed unmyelinated C-low threshold mechanoreceptors marked by Vglut3 expression as a candidate population for driving mechanical hypersensitivity (Seal et al., 2009). However, allodynia persists even when these neurons fail to develop (Lou et al., 2013), and recent evidence indicates that transient Vglut3 expression in spinal interneurons accounts for the phenotype (Peirs et al., 2015). Finally, A-fibers marked by de novo expression of Neuropeptide Y (Ossipov et al., 2002b), TLR5 expression (Xu et al., 2015) or TrkB expression (Peng et al., 2017) have also been suggested to influence mechanical hypersensitivity.

The skin is innervated by an array of functionally distinct populations of mechanoreceptors that can be distinguished by their conduction velocity, response properties, mechanical threshold and the type of end organ that they innervate (Abraira and Ginty, 2013). Molecular markers for many of these populations have recently been identified, amongst which, the receptor tyrosine kinase TrkB, has been proposed to mark both D-hairs (Li et al., 2011) and rapidly adapting Aβ mechanoreceptors (RAMs) (Bourane et al., 2009; Luo et al., 2009; Wende et al., 2012b). These neurons innervate the same hair follicle forming a functional unit (Li et al., 2011), and are amongst the most sensitive of all cutaneous mechanoreceptors, and respond preferentially to dynamic stimuli, making them a strong candidate for the neuronal subtype mediating mechanical allodynia.

Here we generated an inducible TrkB^CreERT2^ mouse line to manipulate this population of neurons in adult mice in vivo. Through loss and gain of function experiments we demonstrate that TrkB positive neurons are both necessary and sufficient for the expression of mechanical hypersensitivity after nerve injury. Because TrkB is a cell surface receptor, we reasoned that its ligand BDNF could allow for pharmacological targeting of these neurons and suppression of their role in driving mechanical allodynia. We therefore generated a recombinant BDNF^SNAP^ fusion protein and conjugated it to the near infrared (IR) photosensitizer IRDye®700DX phthalocyanine (IR700). In vivo delivery of BDNF^SNAP^-IR700 and near IR illumination led to a retraction of TrkB positive sensory neurons from the skin and a prolonged alleviation of mechanical hypersensitivity in multiple models of neuropathic pain. Thus, ligand-guided laser ablation of a subset of mechanoreceptors in the skin is an effective means of selectively silencing the input which drives mechanical allodynia in neuropathic pain states.

## Results

### TrkB positive sensory neurons are D-hair and Rapidly Adapting mechanoreceptors

We generated TrkB^CreERT2^ mice (Figure S1) and crossed them with Rosa26^RFP^ reporter mice to examine colocalization of TrkB with established cellular markers in adult mouse sensory ganglia. Approximately 10% of neurons in the dorsal root ganglia (DRG) were positive for TrkB^CreERT2^, corresponding to the ~8% of cells which expressed TrkB mRNA (Figures 1D and S2). Expression was evident in large neurons marked by NF200, NF200 plus Ret, and TLR5 (Figures S3A-C and 1D), and not present in nociceptors positive for IB4 or CGRP, or C low threshold mechanoreceptors marked by TH (Figures S3A-C and 1D). To assay TrkB^CreERT2^ positive sensory input into the spinal cord we generated a reporter line in which Cre dependent expression of mCherry was driven from the sensory neuron specific Avil locus (Figures S1B and S1C). TrkB^CreERT2^::Avil^mCherry^ positive sensory neurons were present in laminae III/IV of the dorsal horn of the spinal cord where they formed column-like structures extending dorsally (Figure 1I) (Li et al., 2011). We further investigated the projections of TrkB neurons to the skin. TrkB^CreERT2^ fibers formed longitudinal lanceolate endings around hair follicles (Figure 1J) and extended to Meissner corpuscles in the glabrous skin (Figure 1K). We also examined expression of TrkB in human tissue using a TrkB antibody. In agreement with mouse data, TrkB immunoreactivity was present in human DRG in large neurons co-expressing NF200, Ret, and TLR5 but largely absent from nociceptors expressing TrkA (Figures 1E-H and S3D). Similarly, in glabrous skin, TrkB immunoreactivity was detected in NF200 positive fibers innervating Meissner corpuscles (Figure 1L). Collectively, these data indicate that TrkB^CreERT2^ marks a population of putative mechanoreceptive neurons in mouse and human.

**Figure 1:**
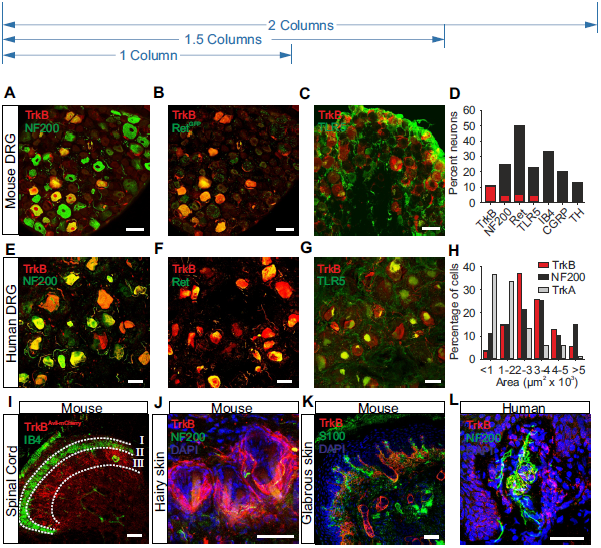
TrkB positive sensory neurons are putative mechanoreceptors. **(A-D)** Double immunofluorescence of DRG sections from TrkB^CreERT2^::Rosa26^RFP^ mice with **(A)** NF200, **(B)** Ret, visualized using TrkB^CreERT2^::Rosa26^RFP^::Ret^EGFP^ triple transgenic mice, **(C)** TLR5, **(D)** Quantification of staining on mouse DRG sections; TrkB^+^ cells account for ~10% of all DRG neurons and all co-express NF200 or NF200+Ret^eGFP^, while they are negative for IB4, CGRP and TH. **(E-H)** Double immunofluorescence of human DRG sections stained with antibodies against TrkB and **(E)** NF200, **(F)** Ret, **(G)** TLR5. **(H)** Size distribution for human DRG neurons expressing TrkB, NF200 and TrkA. **(I)** Section through the lumbar spinal cord of TrkB^CreERT2^::Avil^mCherry^ mice stained with IB4. **(J)** TrkB^+^ lanceolate endings in a section of the back hairy skin of TrkB^CreERT2^::Rosa26^SnapCaaX^ labeled with Snap Cell TMRstar (red), NF200 (green) and DAPI (blue). **(K)** Section from the glabrous skin of TrkB^CreERT2^::Rosa26^ChR2YFP^ (red) stained with anti-S100 a marker for Meissner's corpuscles (green) and DAPI (blue) showing TrkB^+^ innervation. **(L)** Section from human glabrous skin stained with antibodies against TrkB (red) and NF200 (green), and DAPI (blue). Scale bars, A-C and I 50μm, E-G and J-L 40μm. Error bars represent SEM.

To unequivocally establish the identity of TrkB^CreERT2^ positive sensory neurons we characterized their response properties utilizing a combination of electrophysiology and optogenetic activation. Mice expressing the light-gated ion channel channelrhodopsin-2 (ChR2) in TrkB positive cells were generated (TrkB^CreERT2^::Rosa26^ChR2^) and an ex vivo skin nerve preparation was used to identify neuronal subtypes which could be concomitantly activated by light. Strikingly, we determined that all D-hair and RAMs could be stimulated by light whereas all other subtypes of sensory neurons were not responsive (Figures 2A-C). Thus TrkB marks myelinated neurons that innervate hair follicles and Meissner’s corpuscles and are tuned to detect gentle mechanical stimuli.

**Figure 2:**
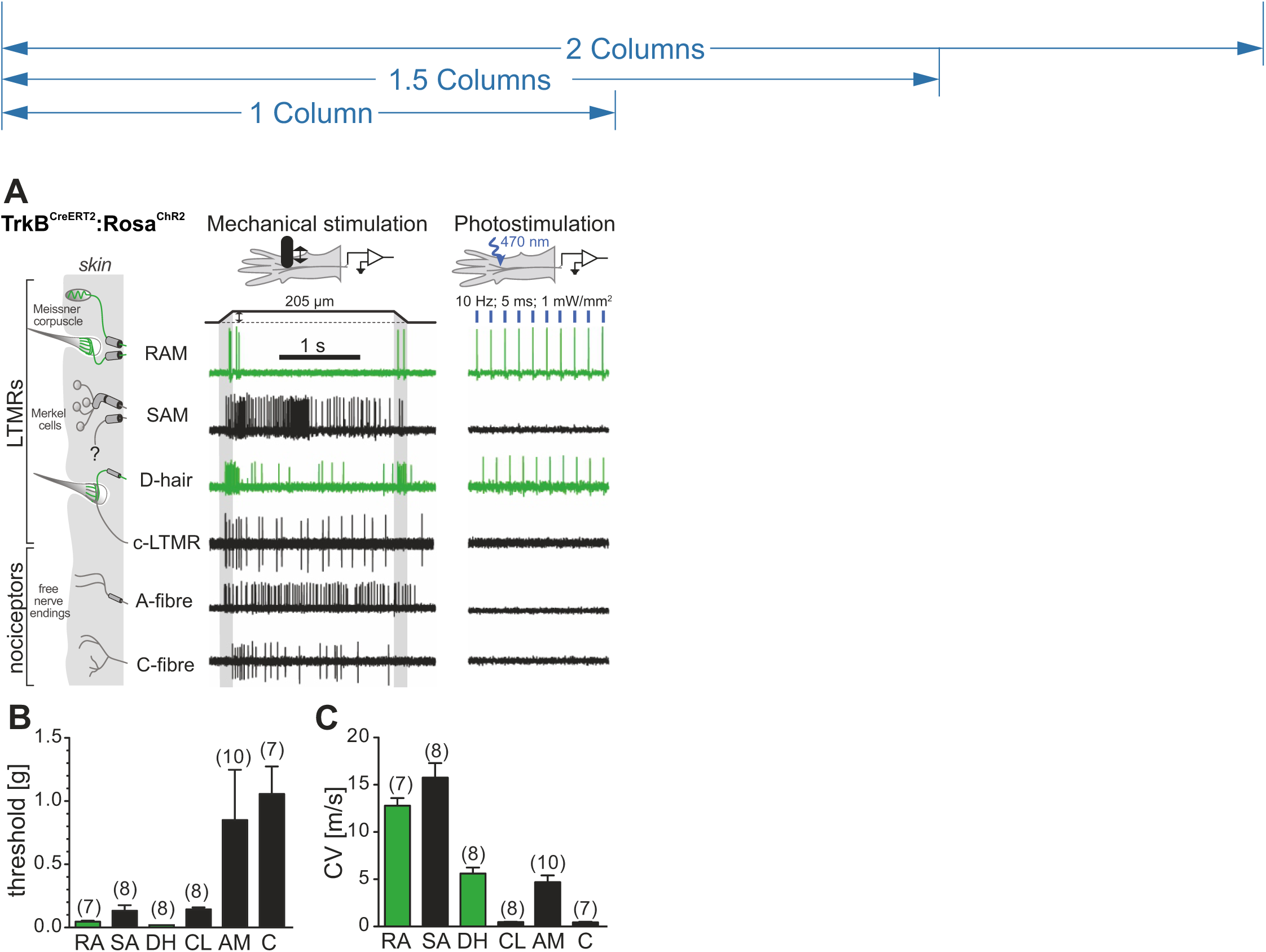
TrkB positive sensory neurons are myelinated low threshold mechanoreceptors. In-vitro skin nerve preparation from TrkB^CreERT2^::Rosa26^ChR2^ mice showing **(A)** representative responses to 10Hz stimulation with blue light, **(B)** the minimal force required to elicit an action potential in the indicated fibre type and **(C)** the conduction velocities of the fibre types. Red bar represents TrkB^+^ afferents, n number indicated in brackets. Error bars represent SEM.

### TrkB positive sensory neurons detect light touch under basal conditions

To determine the role played by TrkB^CreERT2^ positive D-hairs and RAMs in sensory evoked behavior, we genetically ablated TrkB neurons in the peripheral nervous system. We used a Cre-dependent diphtheria toxin receptor driven from the Avil locus (Arcourt et al., 2017; Stantcheva et al., 2016) that allowed for selective deletion of TrkB positive neurons only in adult sensory ganglia. Upon systemic injection of diphtheria toxin we achieved a ~90% ablation of TrkB^CreERT2^::Avil^iDTR^ and TrkB mRNA positive neurons with a parallel reduction in the number of NF200 positive neurons by ~10% and no change in the expression of other markers (Figure 3A-C, Figure S2). We performed a series of behavioral tests in these animals examining sensory responses to a range of thermal and mechanical stimuli applied to the glabrous and hairy skin. There were no differences in responses to evaporative cooling evoked by acetone application (Figures 3D and 3E), or in thresholds to noxious heat (Figures 3F and 3G) after diphtheria toxin ablation. Similarly, grip strength (Figure 3H) was unaltered by ablation of TrkB^CreERT2^ neurons, as were responses to noxious pinprick (Figure 3I), and static or punctate mechanical stimulation of the hairy skin of the back (Figures 3J and 3K). We further examined responses to dynamic mechanical stimuli by monitoring responses to brushing of the plantar surface of the paw. Using a puffed out cotton swab which exerts forces in the range 0.7-1.6mN, we observed a significant reduction in responsiveness upon ablation of TrkB positive neurons (Figure 3L). Intriguingly, these differences were not apparent upon application of stronger dynamic forces using a paint brush (>4mN, Figure 4B) Thus under basal conditions, TrkB positive sensory neurons are required for behavioral responses to the lightest of mechanical stimuli.

**Figure 3:**
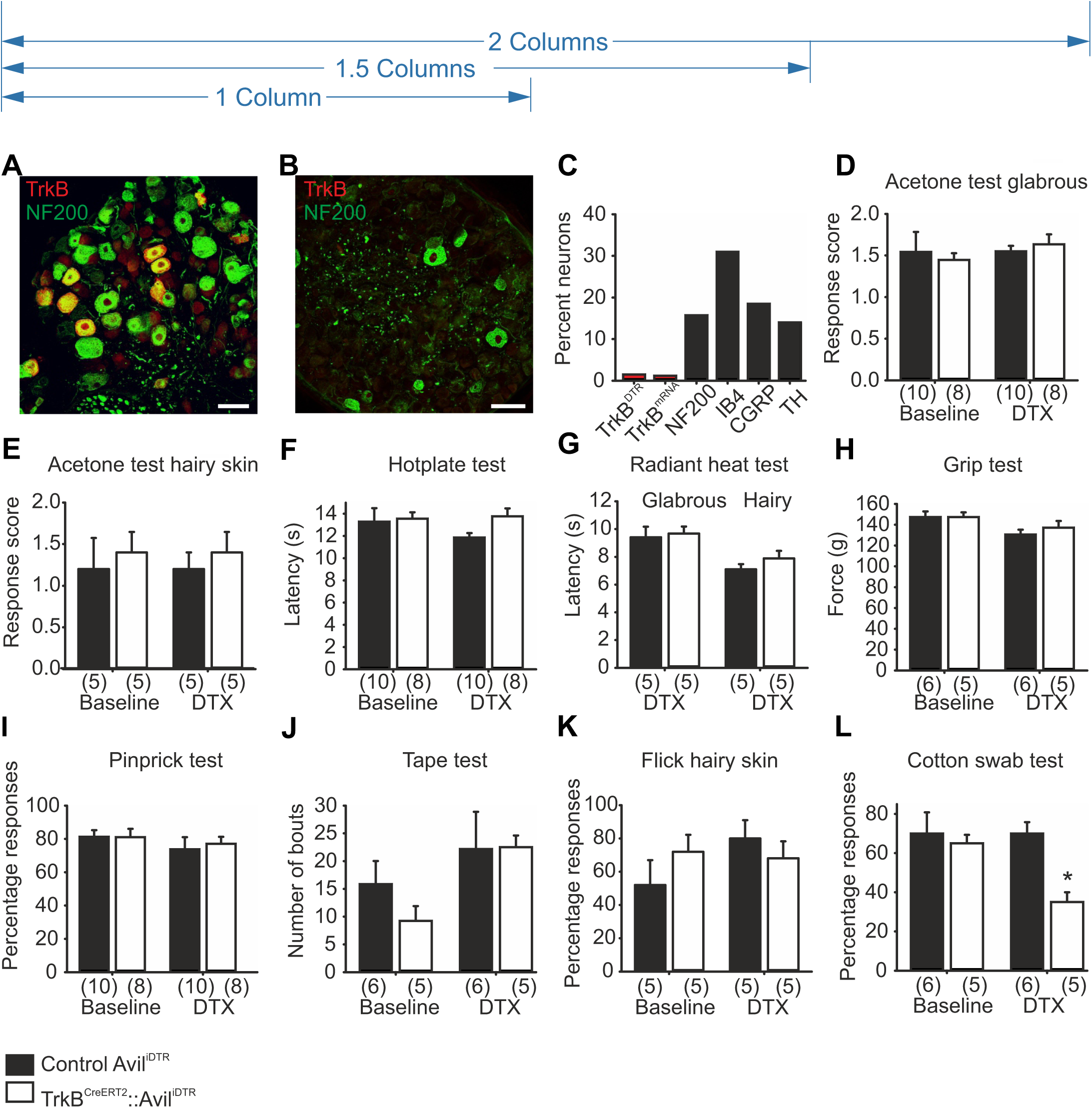
Diphtheria toxin mediated ablation of TrkB+ sensory neurons. Immunostaining of DRG sections of TrkB^CreERT2^::Avil^iDTR^ mice with an antibody against the diphtheria toxin receptor (red) from **(A)** untreated mice and **(B)** after i.p injections of diphtheria toxin. **(C)** Quantification of DRG sections indicating a ~90% decrease in TrkB^DTR^ and TrkB^mRNA^ cells after ablation and ~10% reduction in NF200^+^ neurons without affecting other subpopulations. **(D-L)** Behavioral responses in littermate control mice (Avil^iDTR^, black bars) and TrkB^CreERT2^::Avil^iDTR^ mice (white bars) showing no differences in responses before and after ablation in the acetone drop test (t-test; p>0.05) on **(D)** glabrous and **(E)** hairy skin, **(F)** hot plate test (t-test; p>0.05), **(G)** radiant heat test on glabrous and hairy skin (t-test; p>0.05), **(H)** grip test (t-test; p>0.05), **(I)** pin-prick test (t-test; p>0.05), **(J)** tape test (t-test; p>0.05), and **(K)** punctate mechanical flick test applied to the hairy back skin (t-test; p>0.05). **(L)** Ablated mice show a reduction in sensitivities to cotton swab (t-test, p<0.001). Scale bars in A, B 50μm, error bars indicate SEM.

**Figure 4:**
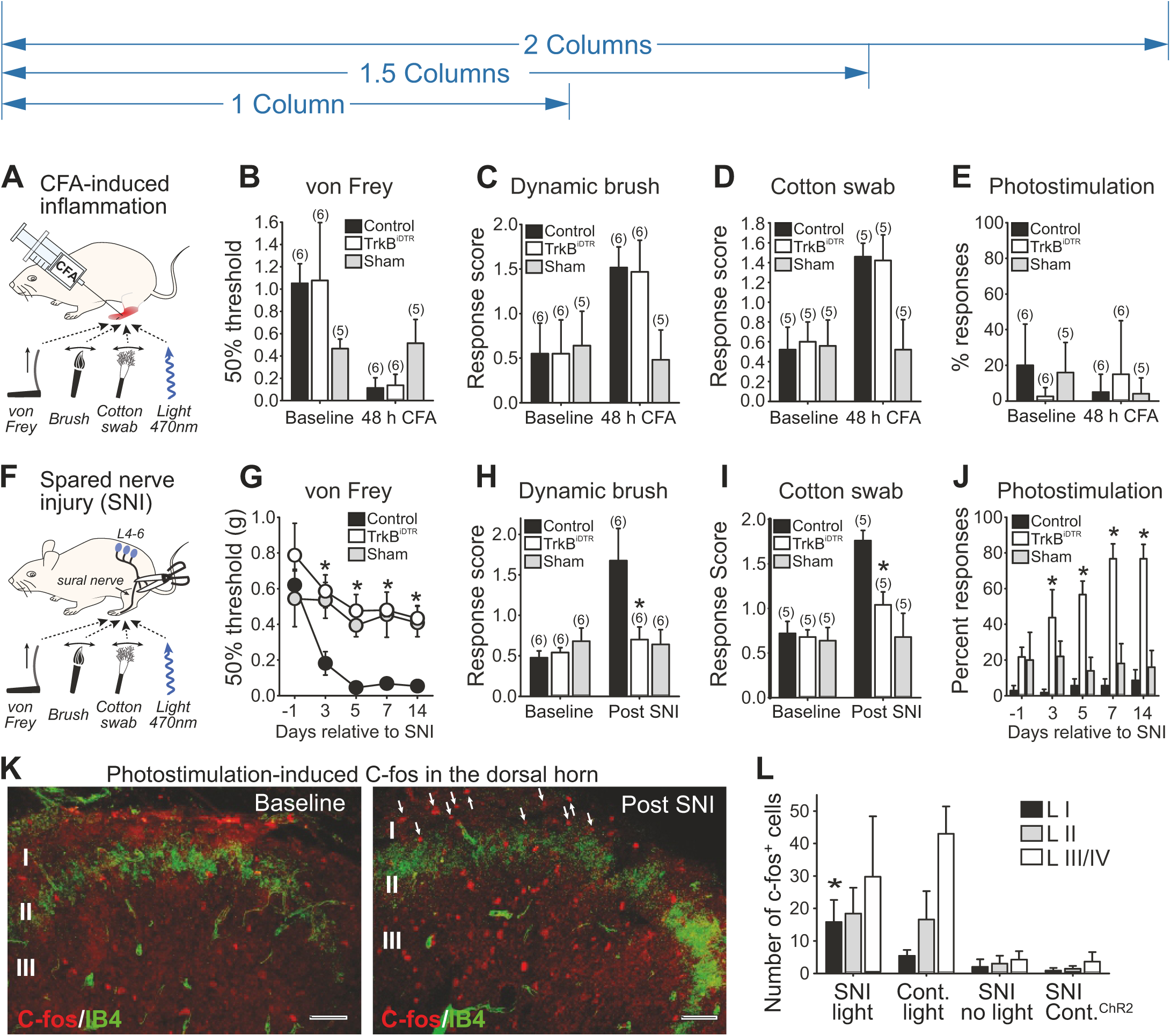
TrkB+ neurons are necessary and sufficient to convey mechanical allodynia after nerve injury. **(A)** Schematic of CFA injection and behavior tests following ablation of TrkB+ neurons. Mechanical hypersensitivity in control Avil^iDTR^ (black bar), TrkB^CreERT2^::Avil^iDTR^ (white bar) and sham injected (grey bar) mice 48 hours after CFA injections as measured by **(B)** von Frey filaments (t-test, p>0.05), **(C)** dynamic brush stimuli (t-test; p>0.05) and **(D)** cotton swab stimuli (t-test; p>0.05). All mice received two diphtheria toxin injections 7 days and 10 days before CFA treatment. **(E)** Paw withdrawal frequencies in control (black bar), CFA injected (white bar) and sham (grey bar) injected paw of TrkB^CreERT2^::Rosa26^ChR2^ mice upon stimulation with 473nm blue light. No significant differences under baseline conditions and 48 hours after CFA injection (Mann-Whitney test; p>0.05). **(F)** Schematic of SNI and behavioral tests following ablation of TrkB+ neurons. **(G)** von Frey mechanical thresholds indicating that ablation of TrkB+ neurons abolished the development of mechanical allodynia after SNI in TrkB^CreERT2^::Avil^iDTR^ mice (white circles) as compared to Avil^iDTR^ controls (black circles) (n=7 for both sets, Two-way RM ANOVA; p<0.001 followed by a Bonferroni post-hoc test). Sham operated mice (grey circles) did not develop mechanical hypersensitivity. **(H-I)** Reduced dynamic allodynia in ablated TrkB^CreERT2^::Avil^iDTR^ mice (white bar) as compared to littermate controls (black bar; t-test p<0.05) stimulated with a **(H)** brush or **(I)** cotton swab. Sham operated mice did not develop dynamic allodynia. **(J)** Nociceptive behavior evoked by optogenetic stimulation of the paws of TrkB^+/+^::Rosa26^ChR2^ (black bars) and TrkB^CreERT2^::Rosa26^ChR2^ (white bars) mice after SNI, or ipsilateral paws (grey bars) of sham operated mice (Two-way RM ANOVA; p<0.001). **(K)** Cross sections of lumbar spinal cord from TrkB^CreERT2^::Rosa26^ChR2^ mice labelled for c-fos (red) and IB4 (green) after 1 minute exposure to 15Hz blue light. Representation of section taken from an uninjured mouse and a section from an injured mouse at 7 days post SNI. **(L)** Quantification of the number of c-fos positive cells in laminae I, II and III/V of the lumbar spinal cord within a 40μm section. Data are shown for SNI, non-injured and sham operated TrkB^CreERT2^::Rosa26^ChR2^ mice, and control SNI Rosa26^ChR2^ mice. Baseline indicates pre-ablation and pre-treatment. Error bars indicate SEM. Scale bars in k 40μm.

### TrkB positive sensory neurons are both necessary and sufficient to produce pain from light touch after nerve injury

On account of the exquisite sensitivity of TrkB positive neurons, we next asked whether they contribute to mechanical hypersensitivity in models of injury-induced pain. We took both a loss of function approach using genetic ablation, and a gain of function approach using optogenetic activation of TrkB neurons. We first considered a model of inflammatory pain by injecting Complete Freund’s Adjuvant (CFA) into the plantar surface of the paw, and monitoring responses to von Frey filaments and dynamic brush or cotton swab stimuli (Figure 4A). Ablation of TrkB neurons in TrkB^CreERT2^::Avil^iDTR^ mice had no effect on any of these measures of mechanical hypersensitivity after inflammation (Figures 4B-D). We next induced neuropathic pain in mice using the Spared Nerve Injury (SNI) model (Figure 4F). Control mice developed a profound mechanical and cold hypersensitivity in the sural nerve territory of the paw (Figures 4G-I and S4). Strikingly, upon ablation of TrkB^CreERT2^::Avil^iDTR^ sensory neurons, mice did not develop mechanical allodynia to either punctate or brushing stimuli (Figures 4G-I), while response to cold stimuli were unchanged (Figure S4). We further examined whether optogenetic activation of TrkB neurons could evoke pain behavior. Using photo-stimulation parameters which evoked robust firing in the ex vivo skin nerve preparation, we observed no discernible behavioral response to light application to the paw either in basal conditions or after CFA-induced inflammation in TrkB^CreERT2^::Rosa26^ChR2^ mice (Figure 4E, and Movie S1). Identical stimulation conditions applied to the hairy skin of the ear auricle evoked a brief ear twitch in TrkB^CreERT2^::Rosa26^ChR2^ mice (Movie S2), likely reflecting activation of the dense network of mechanoreceptors in this structure (Schoebl, 1871). We performed further experiments in mice with the SNI model of neuropathic pain. Three days after injury we observed that selective stimulation of TrkB neurons with light evoked nocifensive behavior. This was evident as a prolonged paw withdrawal from the stimulation, lifting of the paw and licking of the illuminated area (Figure 4J and Movie S3) that continued for several minutes after light application. Such behavior persisted throughout the 2 weeks observation period and was never observed in control mice (Figure 4J). Thus under neuropathic pain conditions, TrkB sensory neurons are necessary and sufficient to convey the light touch signal that evokes pain.

As a neuronal correlate of this apparent pain behavior, we examined induction of the immediate early gene C-fos in the dorsal horn of the spinal cord (Figures 4K-L and Figures S5A-B). In TrkB^CreERT2^::Rosa26^ChR2^ mice without injury, optical stimulation evoked C-fos immunoreactivity primarily in laminae III and IV of the spinal cord, the region where TrkB neurons terminate (Figures 4K and 4L). Upon nerve injury however, identical stimulation parameters induced C-fos staining in lamina I of the dorsal horn (Figures 4K and 4L), an area associated with nociceptive processing. To determine whether this resulted from aberrant sprouting of TrkB+ afferents into superficial laminae after nerve lesion, we examined TrkB+ sensory input into the spinal cord using TrkB^CreERT2^::Avil^mCherry^ mice (Figure S5C and S5D). We were unable to detect any difference in TrkB distribution in mice with SNI, suggesting that de novo expression of C-fos likely arises from plasticity within the interneuron network of the dorsal horn (Cheng et al., 2017; Duan et al., 2014; Foster et al., 2015; Peirs et al., 2015) and not from sprouting (Hughes et al., 2003; Woolf et al., 1992) of TrkB+ mechanoreceptors into the superficial dorsal horn.

### Ligand guided laser ablation TrkB positive sensory neurons

In light of the clinical importance of mechanical allodynia in neuropathic pain patients, we sought to develop a pharmacological strategy to exploit the striking selectivity of TrkB to the peripheral neurons which provoke this pain state. We reasoned that BDNF, the ligand for TrkB, may give access to these neurons and allow for their manipulation in wildtype, non-transgenic animals. To this end we produced recombinant BDNF protein with a SNAP-tag fused to its C-terminus that would enable its chemical derivatization. BDNF^SNAP^ was labelled in vitro with fluorescent SNAP-Surface647 substrate and applied to HEK293T cells expressing neurotrophin receptors. Fluorescently labelled BDNF^SNAP^ displayed remarkable selectivity for its cognate receptor complex TrkB/p75, and did not bind to cells expressing related neurotrophin receptors TrkA/p75 or TrkC/p75 (Figures S6A-C). We further tested whether BDNF^SNAP^ would recognize native TrkB receptors in DRG neurons. BDNF^SNAP^ was conjugated to Qdot 655 quantum dots and applied to dissociated DRG from TrkB^CreERT2^::Rosa26^RFP^ mice. We observed a >95% overlap between BDNF^SNAP^ and TrkB^CreERT2^ positive cells (Figure 5A) indicating that recombinant BDNF^SNAP^ is a highly selective means of targeting TrkB neurons.

**Figure 5:**
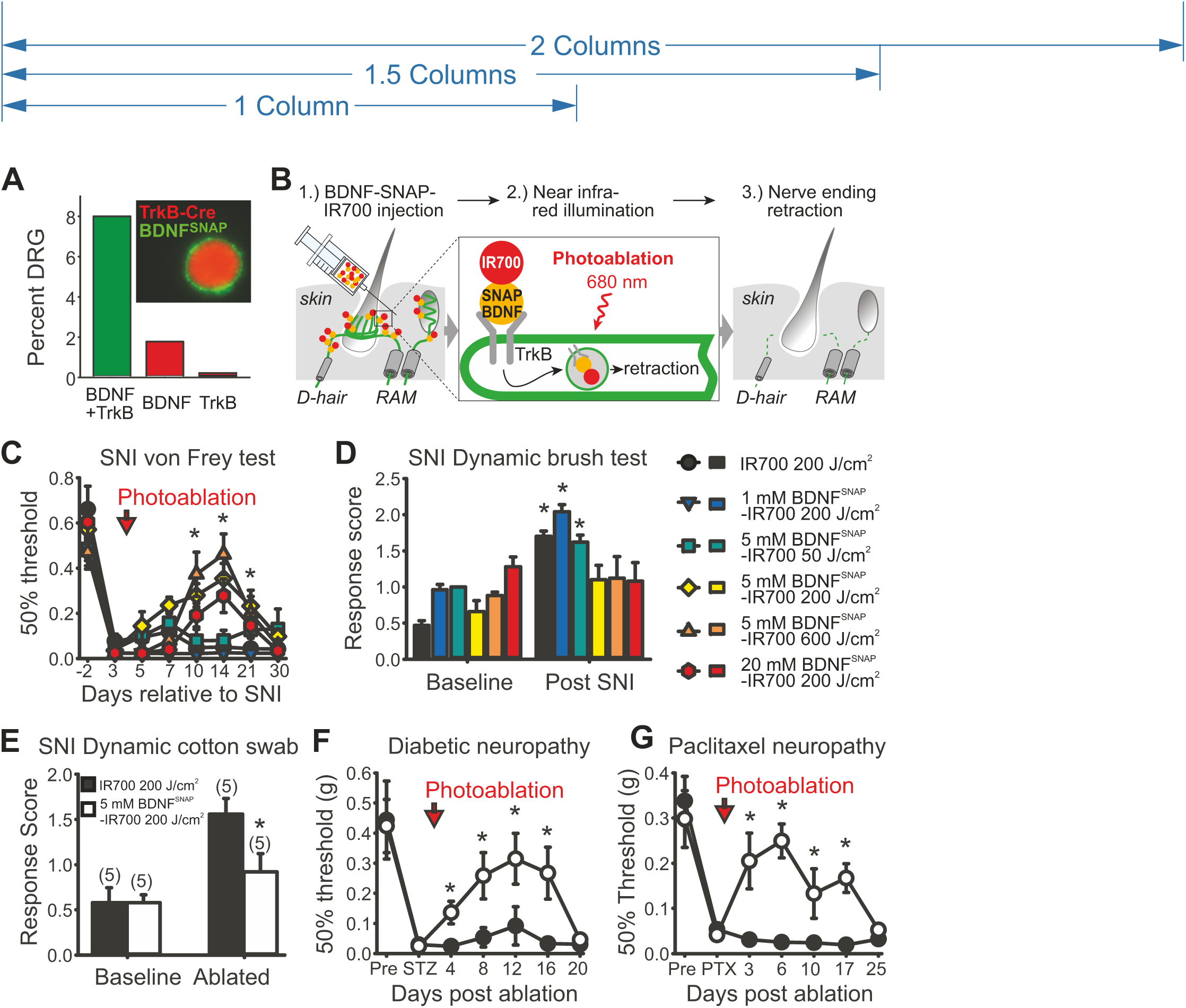
Optopharmacological targeting of TrkB+ neurons with BDNF^SNAP^. **(A)** Labeling (inset) and quantification of dissociated DRG from TrkB^CreERT2^::Rosa26^RFP^ mice with BDNF^SNAP^ shows substantial overlap of BDNF^SNAP^ binding to TrkB+ cells (n=4, 450 cells). **(B)** Schematic representation of BDNF^SNAP^-IR700 injection and photoablation. **(C)** BDNF^SNAP^-IR700 mediated photoablation of the paw of SNI mice results in a dose dependent reversal of mechanical hypersensitivity as assayed with von Frey filaments (n=10, two-way RM ANOVA; p<0.05 followed by a Bonferroni post-hoc test) and **(D)** dynamic brush stimuli (t-test; p<0.05). (**E**) Hypersensitivity to cotton swab is also reversed by photoablation (t-test: p<0.05). **(F)** BDNF^SNAP^-IR700 mediated photoablation reverses mechanical allodynia in the streptozotocin (STZ) model of diabetic neuropathy (n=5, two-way RM ANOVA; p<0.05 followed by a Bonferroni post-hoc test. Open circles; 5μM BDNF^SNAP^-IR700 at 200J/cm^2^, closed circles, 5μM IR700 at 200J/cm^2^). **(G)** BDNF^SNAP^-IR700 mediated photoablation reverses mechanical allodynia in the paclitaxel (PTX) model of chemotherapy induced neuropathy (n=5, two-way RM ANOVA; p<0.05 followed by a Bonferroni post-hoc test. Open circles; 5μM BDNF^SNAP^-IR700 at 200J/cm^2^, closed circles, 5μM IR700 at 200J/cm^2^). Error bars indicate SEM.

To manipulate TrkB neurons in vivo, we reasoned that BDNF^SNAP^ may allow for targeted photoablation of these neurons through delivery of a photosensitizing agent (Mitsunaga et al., 2011 Yang et al., 2015). We synthesized a benzylguanine modified derivative of the highly potent near-infrared photosensitizer IRDye®700DX phthalocyanine (IR700) and conjugated it in vitro to BDNF^SNAP^. In initial experiments we applied BDNF^SNAP^-IR700 to HEK293T cells expressing TrkB/p75 and assayed cell death following near infrared illumination. In cells expressing TrkB/p75 we observed substantial cell death 24 hours after brief illumination that was not evident upon mock transfection or treatment with IR700 alone (Figures S6D-F). We next sought to assess the therapeutic potential of this approach by investigating the effects of BDNF^SNAP^-IR700 mediated photoablation in wildtype mice with neuropathic pain. Upon establishment of robust mechanical allodynia three days after SNI, we injected a range of concentrations of BDNF^SNAP^-IR700 into the ipsilateral paw of injured mice and illuminated the skin with different light intensities (Figure 5B). Strikingly, we observed a concentration and illumination dependent rescue of both von Frey withdrawal thresholds (Figure 5C) and dynamic brush or cotton swab evoked allodynia (Figure 5D and E) that persisted for more than 2 weeks after a single treatment regime. We examined whether such pronounced effects were also evident in other types of neuropathic pain. Indeed, in both the streptozotocin model of painful diabetic neuropathy (Like and Rossini, 1976), and the paclitaxel model of chemotherapy induced neuropathic pain (Apfel et al., 1991), we observed a marked reversal of mechanical hypersensitivity that peaked around 10 days post treatment and returned to injury levels by day 20 (Figures 5F and 5G). To determine the selectivity of this approach, we further assessed the effects of BDNF^SNAP^-IR700 mediated photoablation on behavioral responses under basal conditions. We observed no deficits in sensitivity to cold, heat, or pinprick upon treatment (Figures S7A-C). Responses to cotton swab were also unaffected by photoablation (Figure S7D), perhaps because the skin area that is stimulated in this test (50 mm^2^) extends beyond the zone of illumination (15-20 mm^2^).

### Mechanism of BDNF^SNAP^-IR700 mediated reversal of mechanical hypersensitivity

Using a TrkB^CreERT2^::Rosa26^SNAPCaaX^ reporter mouse line (Yang et al., 2015) to identify TrkB positive afferents, and a PGP9.5 antibody to label all fibers, we examined the innervation density of hypersensitive skin over the course of phototherapy. Prior to photoablation, we detected TrkB positive lanceolate endings around hair follicles (Figure 6A) and innervation of Meissner corpuscles in the plantar surface of the paw (Figure S8E). At 7 days after photoablation (13 days post-SNI) when behavioral reversal of mechanical hypersensitivity was most pronounced, we observed selective loss of TrkB fibers but persistent innervation by PGP9.5 fibers in hairy and glabrous skin (Figures 6B, 6F and S8). Indeed, many hair follicles displayed a complete loss of TrkB innervation but still contained PGP9.5 positive circumferential and free nerve endings demonstrating the remarkable specificity of ablation (Figure S9). At 24 days post-photoablation when mechanical hypersensitivity had reverted, TrkB positive fibers were again seen innervating their appropriate end organs in both glabrous and hairy skin (Figures 6C and S8). Importantly, we observed no apparent reduction in innervation of control tissue injected with unconjugated IR700 and illuminated (Figure S9). We further investigated whether loss of TrkB^CreERT2^ neurons was also evident at the level of the cell soma by analyzing the number of TrkB^CreERT2^ positive neurons in the DRG. We observed no difference in the proportion of TrkB neurons 10 days after photoablation (FigureS 6D-F) indicating that the loss of fibers likely reflects local retraction from their peripheral targets.

**Figure 6:**
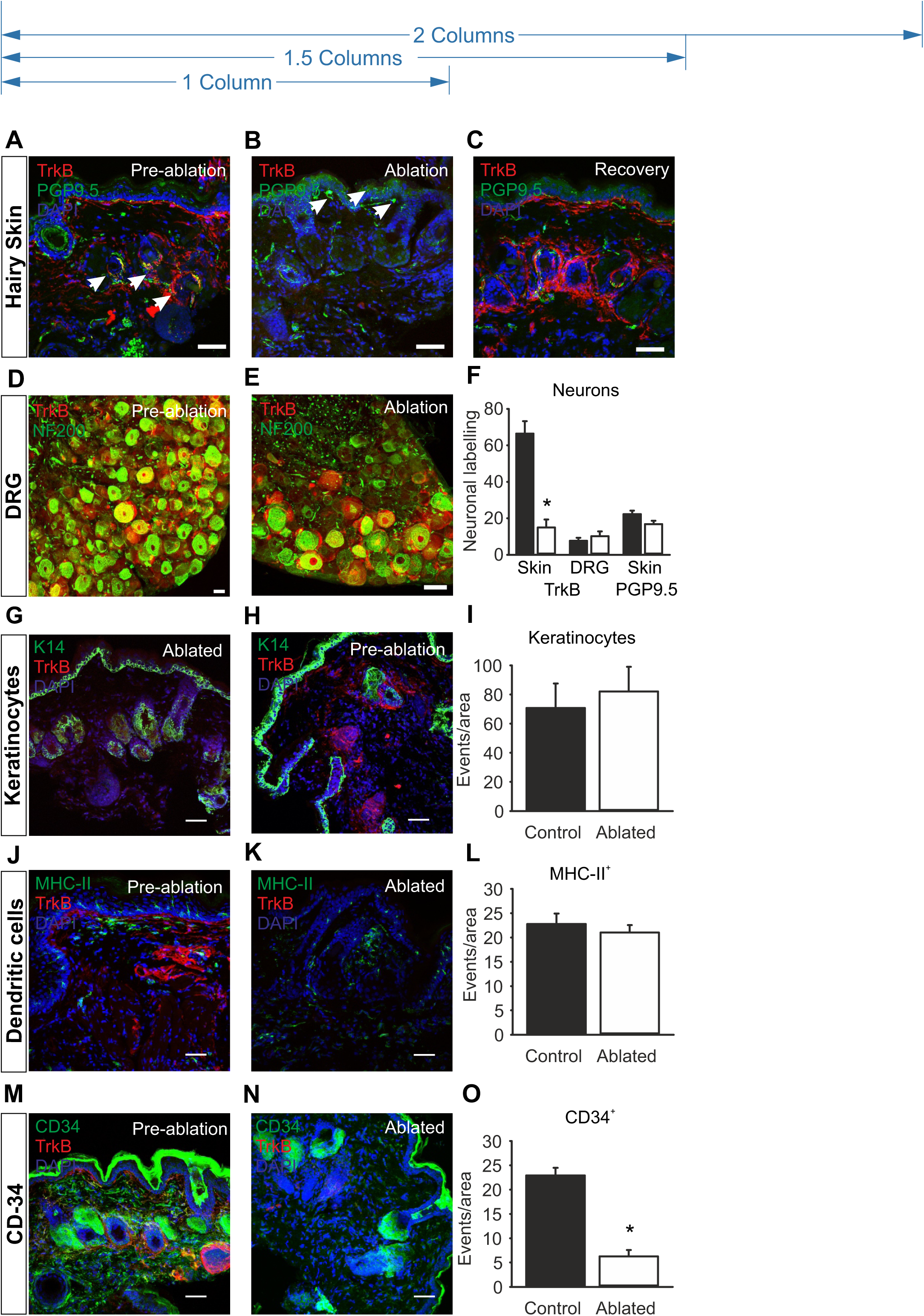
BDNF^SNAP^-IR700 photoablation promotes local retraction of TrkB+ afferents. **(A-C)** Substantial loss of TrkB^CreERT2^ positive afferents (red), but persistence of other fibers (green) upon BDNF^SNAP^-IR700 mediated photoablation. **(A)** Innervation of paw hairy skin prior to ablation, arrows show lanceolate endings. **(B)** Loss of TrkB^CreERT2^ afferents after ablation, arrows show PGP9.5 fibers. **(C)** Reinnervation of skin by TrkB^CreERT2^ afferents at 24 days post ablation. **(D)** DRG section from control TrkB^CreERT2^ mouse labelled for RFP (red) and NF200 (green). **(E)** DRG section from photoablated TrkB^CreERT2^ mouse labelled for RFP (red) and NF200 (green). **(F)** Quantification of the proportion of hair follicle innervation and DRG neurons positive for TrkB following photoablation in the paw and the quantification of PGP9.5+ free nerve endings showing the numbers of free-nerves remain unaffected. Representative skin sections from control and BDNF^SNAP^-IR700 photo-ablated mice labelled with the indicated antibodies. TrkB positive cells are indicated in red and DAPI positive nuclei in blue. **(G-H)** Keratinocytes labelled with K14 (green). **(I)** Quantification of the number of K14+ cells. **(J-K)** Dendritic cells and dermal antigen presenting cells labelled with MHC-II (green) and **(L)** the quantification for MHC-II+ cells. **(M-N)** Mast cells and epithelial and endothelial progenitor cells labelled with CD34 (green). **(O)** Quantification of the number of K14+ cells. Scale bars 40μm. Error bars indicate SEM.

TrkB is also expressed by other cells in the skin in addition to sensory fibers (Botchkarev et al., 2006; Ichikawa et al., 2001; Peng et al., 2013; Truzzi et al., 2011). We sought to identify these cell types and determine whether they are lost upon photoablation and contribute to the behavioral phenotype. TrkB was not detected in Merkel cells, keratinocytes, or dendritic and dermal antigen presenting cells (Figures S10, 6G and 6J), and BDNF^SNAP^-IR700 mediated photoablation did not alter their numbers in the skin (Figures 6H, K and 6I, L). Expression of TrkB was however evident in cells labelled with CD34, a marker of mast cells and epithelial and endothelial progenitor cells (Figure 6M). Moreover, photoablation significantly reduced the number of CD34 positive cells in the skin (Figures 6N-O).

To determine whether it is loss of CD34+ cells or TrkB+ afferents which influences sensory behavior, we injected BDNF^SNAP^-IR700 into the sciatic nerve at mid-thigh level of SNI mice and illuminated the nerve to ablate TrkB sensory fibers but spare CD34 cells in the skin. We first examined histological indicators at the site of injection. In control samples, TrkB+ fibers were clearly visible in both longitudinal and cross sections of nerve, whereas after illumination they were essentially eliminated (Figures 7A-D and G). Importantly we did not detect any CD34+ cells in nerve samples, either before or after illumination (Figures 7A-D). Similarly, TrkB expression was also not evident in other non-neuronal cell type in the sciatic nerve such as S100 positive Schwann cells (Figures 7E and 7F). Finally, we quantified the number of DAPI positive nuclei in nerve sections as a measure of non-neuronal cell density. There was no overlap between DAPI and TrkB expression (Figure 7A, C, and E-G) indicating that TrkB is restricted to neurons in the nerve; indeed we observed an increase in DAPI positive nuclei after illumination, likely reflecting immune cell infiltration upon photoablation. Collectively, these data indicate that BDNF targeted photoablation selectively and effectively eliminates TrkB positive fibers.

**Figure 7:**
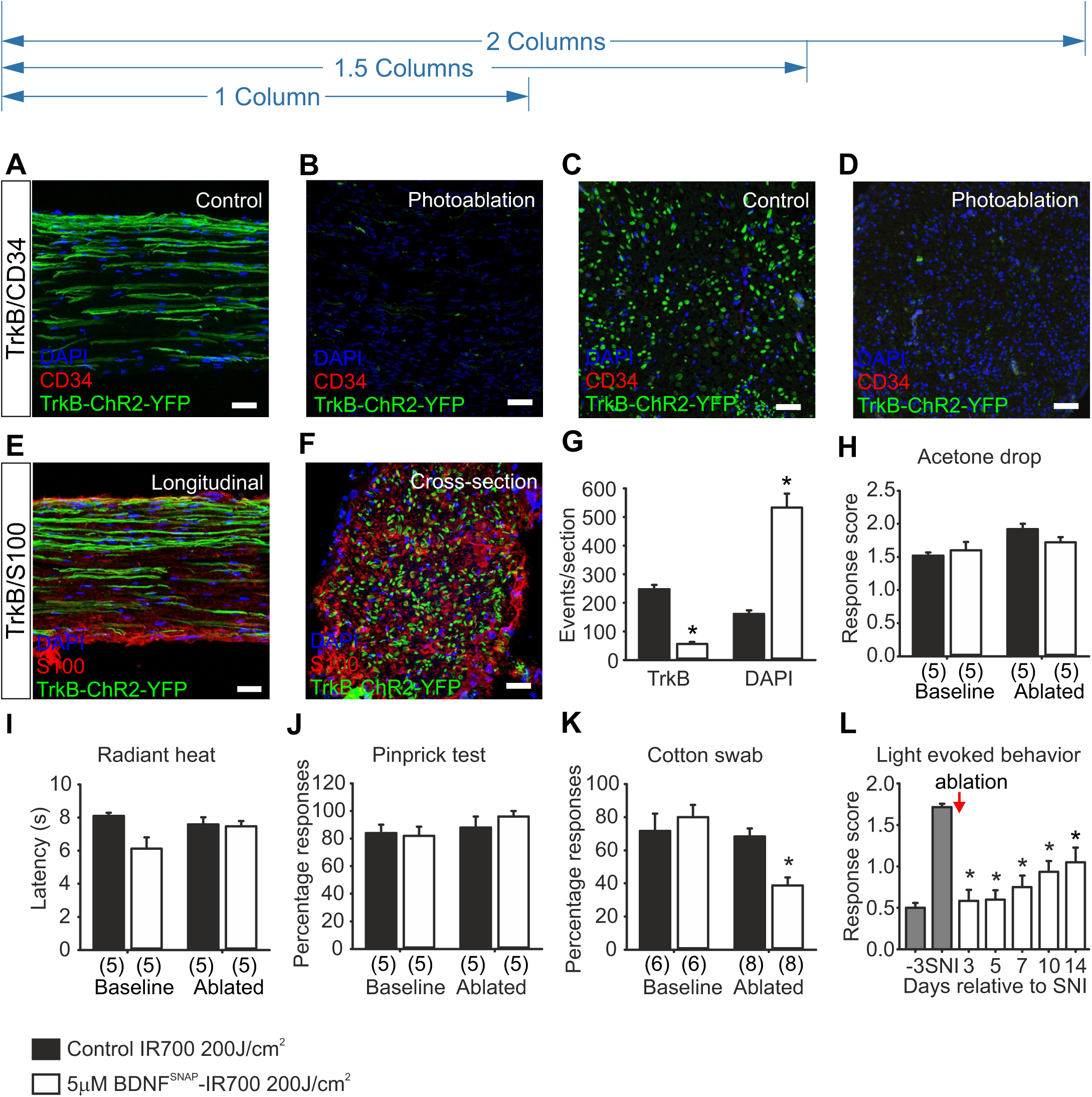
Photoablation of the sciatic nerve shows requirement of TrkB+ neurons in mediating allodynia. **(A-F)** Representative images of the sciatic nerve labelled for TrkB in green, DAPI in blue and CD34 or S100 in red. Longitudinal sections of the nerve show reduction in TrkB+ fibres, but no detectable CD34 in **(A)** non-ablated control and **(B)** photoablated mice. Cross-section of the sciatic nerve from **(C)** control and **(D)** photoablated mice shows reduction in TrkB+ fibres. No co-localization between TrkB+ fibres and S100+ cells in **(E)** longitudinal sections and **(F)** crosssection of the nerve. **(G)** Quantification of the numbers of TrkB+ fibers and DAPI labelled cells in cross sections of sciatic nerve. Behavioral sensitivity following BDNF^SNAP^-IR700 mediated ablation in the sciatic nerve: **(H)** Acetone drop test (t-test; p>0.05), **(I)** radiant heat test (t-test; p>0.05), and **(J)** pin-prick test (t-test; p>0.05) are not altered by nerve photoablation. However sensitivity to **(K)** cotton swab (t-test; p<0.05) in control animals, and **(L)** light evoked behavior in TrkB^CreERT2^::Rosa26^ChR2^ mice with SNI, are reduced by nerve photoablation (Two-way RM ANOVA; p<0.001). White bars 5μM BDNF^SNAP^-IR700 at 200J/cm^2^, black bars 5μM IR700 at 200J/cm^2^. Baseline indicates pre-ablation and pre-treatment. Error bars indicate SEM. Scale bars a-d 40μm, e 10μm.

We further explored the behavioral consequence of TrkB fiber ablation in the sciatic nerve. In animals which received BDNF^SNAP^-IR700 and illumination of the nerve, behavioral responses to cooling, heating and pinprick were normal (Figures 7H-J). However, sensitivity to cotton swab was significantly reduced (Figure 7K), paralleling the results using genetic ablation of TrkB+ neurons. Finally, we investigated whether optogenetically evoked pain behavior in TrkB^CreERT2^::Rosa26^ChR2^ mice with SNI is also reduced by BDNF^SNAP^-IR700 nerve injection and illumination. Upon photoablation of TrkB+ fibers in the sciatic nerve we observed a significant reduction in light driven nocifensive behavior in TrkB^CreERT2^::Rosa26^ChR2^ mice (Fig. 4L). Thus, TrkB+ sensory afferents, and not CD34+ cells in the skin, likely underlie behavioral sensitivity to light touch under basal conditions and after nerve lesion.

## Discussion

Mechanical allodynia is a cardinal feature of neuropathic pain that is challenging to treat and exerts a substantial societal burden. Here we identify the first relay station in the neuronal pathway that confers pain from gentle touch under neuropathic pain states. We demonstrate that TrkB positive sensory neurons detect the lightest touch under basal conditions but after nerve injury are both necessary and sufficient to drive mechanical allodynia. We further describe new technology based upon ligand-mediated delivery of a phototoxic agent to target these neurons and reverse mechanical hypersensitivity in neuropathic pain states.

### TrkB as a marker for LTMR’s

Our work on the identity of TrkB positive sensory neurons builds upon previous studies that have reported that TrkB is expressed in two populations of myelinated mechanoreceptors that are differentiated by co-expression of Ret, and form longitudinal lanceolate endings around hair follicles (Bourane et al., 2009; Li et al., 2011; Usoskin et al., 2015; Wende et al., 2012a). We demonstrate that TrkB^+^ afferents also innervate Meissner’s corpuscles in the glabrous skin, and that TrkB marks essentially all Aδ-LTMR’s (also known as D-hairs) and RA Aβ-LTMR’s, but no other cutaneous sensory neuron subtype, establishing the TrkB^CreERT2^ mouse line as a powerful tool for investigating the function of these neurons in vivo. Moreover, our histological analysis of human tissue indicates that DRG neurons and skin biopsies of human subjects have comparable molecular profiles to mice, suggesting that TrkB+ neurons in humans may also form LTMRs.

### TrkB^+^ neurons are required to detect gentle mechanical stimuli

To explore the role of Aδ-LTMR’s and RA Aβ–LTMR’s in sensory evoked behavior, we took both a loss of function approach using genetic ablation, and a gain of function approach using optogenetic activation of TrkB+ neurons. We found that TrkB+ neurons were required for a behavioral response to neurons, as were response to the gentlest of dynamic touches, but that their ablation had no influence on responses evoked by thermal or stronger static mechanical stimuli. Similarly, blue-light stimulation of the nape of the neck or the border of the ear in TrkB^CreERT2^::Rosa26^ChR2^ mice elicited a flicking of the ears or the head (Movie S2) that was different from previously described wiping or itching in response to pain (Wilson et al., 2011). It was shown as early as the 1940's that the slightest movement provoked by a cotton-bristle is enough to activate D-hairs, and that this may represent sensations of tickle or gentle blowing of air (Zotterman, 1939). It is intriguing to speculate that optogenetic activation of TrkB+ neurons in the ear may elicit a similar sensation.

Previously, TrkB^+^ D-hairs have been shown to specifically innervate hair follicles in a direction dependent manner and respond to movement of hair in the caudal-to-rostral direction, suggesting that they may be important for tactile determination of direction (Li et al., 2011; Rutlin et al., 2014). We did not determine whether mice had deficits in orientating towards a moving mechanical stimulus in the absence of TrkB+ neurons. However, the ablation approach described here would be valuable for investigating the contribution of these neurons towards orientation. In the same light, development of more intricate tests to measure gentle tactile stimulation of the skin will help better understand the behavioral consequences of stimuli transduced by TrkB^+^ neurons.

### TrkB^+^ neurons drive mechanical allodynia after nerve injury

We further reasoned that manipulation of TrkB^+^ neurons through loss and gain of function experiments would also allow us to identify the peripheral neuron type which inputs mechanical hypersensitivity into the spinal cord. Both nociceptors and LTMRs have been implicated in mechanical allodynia but consensus on the sufficiency and necessity of a specific subpopulation of neurons that conveys this sensation is lacking (Abrahamsen et al., 2008; Ossipov et al., 2002a; Ossipov et al., 2002b; Peng et al., 2017; Seal et al., 2009; Tarpley et al., 2004; Xu et al., 2015). We found that TrkB+ LTMRs were required for hypersensitivity to both punctate and dynamic mechanical stimuli after nerve injury, and that optogenetic activation of TrkB+ neurons was sufficient to evoke strong nocifensive behavior. Importantly, we also determined that in the CFA model of inflammatory pain, TrkB+ neurons were dispensable for mechanical hypersensitivity. Thus, mechanical pain arising from neuropathy or tissue inflammation is likely mechanistically different and should therefore be treated as a distinct clinical entity. Of note, the neurons marked in the TrkB^CreERT2^ line do not express C-fiber markers such as IB4, CGRP and Vglut3, but overlap with markers of A-fibers, all of which have previously been implicated in mechanical allodynia (Abrahamsen et al., 2008; Ossipov et al., 2002a; Ossipov et al., 2002b; Peng et al., 20'7; Seal et al., 2009; Tarpley et al., 2004; Xu et al., 2015). A future challenge will be to determine whether mechanical hypersensitivity is conveyed by only Aδ-LTMR’s or RA Aβ-LTMR’s fibers, or whether both afferent types are required.

We did not explore in detail the mechanistic basis of how TrkB+ neurons provoke mechanical allodynia after nerve injury, focusing instead on developing a potentially translatable tool for its treatment. However our data do give some insights into events in the spinal cord post-injury. For example, using a TrkB^CreERT2^::Avil^mCherry^ mouse line to label TrkB+ afferent projections into the dorsal horn, we detected no gross differences in their termination pattern after nerve injury, arguing against sprouting of LTMR's from lamina III/IV into lamina II (Woolf et al., 1992). We did however observe c-fos labelling in lamina I evoked by optogenetic activation of TrkB+ afferents in the hind-paw of injured mice, indicating that LTMR’s do indeed gain access to pain transmitting neurons in lamina I following nerve injury. Recent evidence suggests that this may arise both through altered gene expression in TrkB neurons (Peng et al., 2017) and through a defect in the feed-forward inhibition of interneurons within laminae III/IV (Braz et al., 2012; Duan et al., 2014; Foster et al., 2015; Torsney and MacDermott, 2006). The TrkB^CreERT2^ line and tools described here will allow for further investigations into these mechanisms.

### BDNF^SNAP^ mediated photo-ablation reverses mechanical allodynia

Current clinical options for reducing neuropathic pain include opioids (like morphine and oxycodone), anti-epileptics such as gabapentin, and tricyclic antidepressants (Attal and Bouhassira, 2015; Dworkin et al., 2010; Dworkin et al., 2007). These drugs have limited effectiveness, serious safety issues, and long term use can lead to addiction (Grosser et al., 2017). Cutaneous receptors in the skin represent an attractive target for novel analgesics, however many potential therapies under development, such as anti-NGF antibodies, and sodium channel blockers are geared towards blocking the function of nociceptors. Our data indicates that LTMRs would be a more appropriate target for alleviating mechanical allodynia, and indeed small molecule inhibitors that silence mechanoreceptors (Wetzel et al., 2017), or inhibit electrical activity in Aβ-LTMR’s (Xu et al., 2015) have been shown to be effective at reducing mechanical allodynia in mouse models. Our approach takes this further by utilizing the TrkB ligand BDNF to deliver a photosensitizer directly to TrkB+ neurons, selectively targeting those neurons which initiate mechanical allodynia. We show that this leads to long term reversal of mechanical hypersensitivity across models of traumatic, diabetic and chemotherapy induced neuropathy, with minimal effects on other sensory modalities.

Application of BDNF^SNAP^-IR700 to the skin and subsequent illumination led to the local retraction of TrkB+ neurons from their end-organs, followed by a re-innervation of appropriate targets 3 weeks later that paralleled the return of mechanical allodynia. This was a remarkably selective process, as illustrated by the continued presence of circumferential endings around hair follicles (likely TrkC/Ret positive field receptors (Bai et al., 2015)), and free nerve endings in the epidermis. Indeed, it is feasible that ligand-targeted photoablation could also be applied to other subtypes of sensory neurons using different ligands to inhibit other sensations. Beyond the therapeutic potential of such an approach, this may also have value as an experimental tool for exploring the consequences of subtype specific nerve ablation and the events that lead to regeneration.

BDNF^SNAP^BDNF^SNAP^-IR700 mediated ablation also allowed us to address a critical question pertaining to the role of TrkB expressing cells in the skin and their contribution to allodynia. For example, it has been demonstrated that optogenetic activation of keratinocytes can trigger action potentials in some populations of cutaneous sensory neurons (but not Aδ-LTMR’s or RA Aβ-LTMRs) and initiate nociceptive behavior (Baumbauer et al., 2015). While we detected no expression of TrkB in keratinocytes, Merkel cells or dendritic cells in the skin, TrkB was observed in mast cells and epithelial and endothelial progenitor cells marked by CD34. Moreover, photoablation reduced the number of CD34^+^ cells in the skin in addition to TrkB^+^ fibers. To determine which of these cell types underlies the behavioral phenotype, we performed photoablation on the sciatic nerve. Importantly, CD34^+^ cells were not evident in the nerve, and photoablation of the nerve produced a phenotype similar to our skin injections, suggesting that CD34^+^ skin cells are not responsible. TrkB has also been reported to be expressed by Schwann cells in the nerve (Frisen et al., 1993), however we were unable to detect overlap of TrkB with the Schwann cell marker S100. Further experiments using a transectional approach to limit ChR2 expression to TrkB fibers, or a triple transgenic TrkB^CreERT2^::Avil^iDTR^::Rosa26^ChR2-YFP^ mouse line with which to perform optogenetic activation in the absence of TrkB^+^ fibres, would further clarify the role of afferent fibers versus other cell types in mediating mechanical allodynia.

In summary, here we identify the peripheral neuronal substrate that confers pain from gentle touch under neuropathic pain states. We demonstrate that TrkB marks a population of sensory neurons that normally detect the lightest touch but drive mechanical hypersensitivity after nerve injury. We further describe new technology based upon a phototoxic derivative of BDNF to target these neurons and reverse allodynia in multiple types of neuropathic pain. This approach is analogous to clinically approved capsaicin patches, in which a high concentration of capsaicin is applied to the skin and leads to retraction of nociceptive fibers (Anand and Bley, 2011). Instead, here we target directly the neurons responsible for mechanical allodynia, allowing for local, on demand treatment of pain through application of light. Further use of this technology and of the genetic tools developed here to manipulate TrkB neurons will now allow precise characterization of the central circuits that gate mechanical pain and transform a normally innocuous sensation into a noxious one.

## Author Contributions

R.D and P.A.H conceived the study. R.D performed the experiments with help from C.M.A, P.P, S.R, L.N. F.J.T and S.G.L performed the electrophysiological studies. C.P, M.M and F.F performed in-vitro analyses with BDNF^SNAP^. F.C.R generated the TrkB^CreERT2^ line. E.P and S.G helped acquiring skin biopsies and with the histological analysis. L.R, S.B, A.F.H and K.J with the production of IR700, R.D and P.A.H wrote the manuscript with essential input from all authors.

## Acknowledgments

We thank Philip Hublitz of EMBL Gene Expression Services, Pedro Moreira of EMBL Transgenic Services, and Violetta Paribeni for technical support of our work. We also acknowledge the assistance of David Hacker, Laurence Durrer and Soraya Quinche of the Protein Expression Core Facility of the EPFL in generation of BDNF^SNAP^. We acknowledge the use of tissues procured by the National Disease Research Interchange (NDRI) with support from NIH grant 2 U42 OD0m11158. This work was funded by EMBL and the Deutsche Forschungsgemeinschaft (SFB H58).

## Supplementary Figure Legends

**Figure S1: Generation of TrkB^CreERT2^ and *Avil^mCherry^ mouse lines.***

**(A)** Schematic representation of ET-recombination based insertion of a Cre^ERT2^ gene cassette into the coding region of a BAC containing the mouse TrkB gene locus **(B)** Schematic diagram of the wild type *Avil* locus with the Avil^hM3Dq-mCherry^ targeting construct, targeted allele and recombination product. **(C)** Southern blot of positive ES clone.

**Figure S2: TrkB mRNA expression in DRG from control and ablated mice.**

**(A)** In situ hybridization showing expression of TrkB mRNA in ~8% of neurons in DRG sections from control mice. **(B)** In situ hybridization showing substantial reduction in TrkB mRNA positive cells following diphtheria toxin mediated ablation. **(C)** Quantification of TrkB mRNA positive neurons in control and ablated mice (t-test; p<0.05). Scale bars, 50μm. Error bars SEM.

**Figure S3. TrkB positive sensory neurons are not expressed in nociceptors.**

Double immunofluorescence of DRG sections from TrkB^CreERT2^::Rosa26^RFP^ mice showing that TrkB does not co-localize **(A)** IB4, **(B)** CGRP, **(C)** TH. **(D)** Section of human DRG showing TrkB is not expressed in TrkA positive neurons.

**Figure S4. Cold allodynia unaffected upon ablation of TrkB+ neurons**

**(A)** Cold hypersensitivity as assayed using the acetone drop test does not differ between TrkB^CreERT2^::Avil^iDTR^ and Avil^iDTR^ control mice (t-test; p>0.05). Error bars SEM.

**Figure S5. C-fos and primary afferent TrkB expression in spinal cord sections.**

**(A)** C-fos expression in spinal cord sections from sham operated TrkB^CreERT2^::Rosa26^ChR2^ mice with light stimulation. **(B)** C-fos expression in spinal cord sections from control Rosa26^ChR2^ mice stimulated with light at 7 days post SNI. **(C and D)** TrkB^+^ primary afferent distribution in TrkB^CreERT2^::Avil^mCherry^ mice at 7 days post SNI. Spinal cord ipsilateral to the injury is shown in **(C)** and contralateral in **(D)**. Scale bars, 40μm.

**Figure S6. BDNF^SNAP^ labelling and IR700 mediated photoablation in vitro.**

**(A-C)** BDNF^SNAP^ labeling of HEK293T cells transfected with **(A)** TrkB/p75NTR, **(B)** TrkA/p75NTR, or **(C)** TrkC/p75NTR. **(D)** Staining of HEK293T cells transfected with TrkB/p75NTR with propidium iodide 24 hours after treatment with BDNF^SNAP^-IR700 and near infrared illumination. **(E)** Staining of mock transfected HEK293T cells with propidium iodide 24 hours after photoablation following treatment with BDNF^SNAP^-IR700. **(F)** Staining of HEK293T cells transfected with TrkB/p75NTR with propidium iodide 24 hours after treatment with IR700 alone and near infrared illumination. Scale bars 50μm.

**Figure S7. BDNF^SNAP^-IR700 mediated photoablation in the paw does not affect baseline sensory behavior**

Responses to **(A)** acetone drop test (t-test; p>0.05), **(B)** hot plate test (t-test; p>0.05), **(C)** pinprick test (t-test; p>0.05) and **(D)** cotton swab test (t-test; p>0.05). White bars 5μM BDNF^SNAP^-IR700 at 200J/cm^2^, black bars 5μM IR700 at 200J/cm^2^. Baseline indicates pre-ablation and pretreatment. Error bars indicate SEM.

**Figure S8. Innervation of the plantar skin of the paw upon BDNF^SNAP^-IR700 mediated photoablation.**

**(A)** Schematic of the mouse paw with boxes showing areas of analysis. 1 indicates hairy skin and is show in Figure 4 m-p. 2. Indicates glabrous skin and is shown here in b-d. 3. Indicates glabrous skin enriched in Meissner corpsucles and is shown in e-g. **(B)** Glabrous skin section prior to photoablation. Note the large number of TrkB^CreERT2^ and PGP9.5 positive fibers that transition through this area. **(C)** Loss of TrkB^CreERT2^ fibers upon ablation. PGP9.5 fibers are still present. **(D)** Recovery of TrkB^CreERT2^ innervation 24 days post ablation. **(E)** Glabrous skin section prior to photoablation are from 3. Note the dense innervation of Meissner corpuscles by TrkB^CreERT2^ fibres. (e) Loss of TrkB^CreERT2^ fibers but not PGP9.5 free nerve endings upon ablation. **(F)** Recovery of TrkB^CreERT2^ fibers 24 days after ablation and reinnervation of Meissner corpuscles. Scale bars 40μm.

**Figure S9. Specificity of BDNF^SNAP^ mediated photoablation**

**(A)** High magnification image of a hair follicle after ablation. Note the absence of TrkB^CreERT2^ fibers (red) but PGP9.5 positive circumferential and longitudinal lanceolate endings (green). **Figure S10. Merkel cells from adult mice are TrkB-Cre negative.**

**(A)** Representative image showing Merkel cells (green) labelled with an antibody again CK20. TrkB staining (red) is not present. Scale bar 40μm.

## Supplementary Movie Legends

**Movie S1: Response to optogenetic activation of the hindpaw after CFA**

Video showing the response of a TrkB^CreERT2^::Rosa26^ChR2^ mouse before and after injection of CFA.

**Movie S2: Response to optogenetic activation on the ear**

Light activation directed towards the ear evokes a flicking response in TrkB^CreERT2^::Rosa26^ChR2^, while control mice do not respond to light.

**Movie S3: Response to optogenetic activation of the hindpaw after SNI**

Light activation evokes strong paw withdrawal 7 days after SNI in TrkB^CreERT2^::Rosa26^ChR2^ mice, while under baseline conditions, mice do not respond to light.

